# Castanet: a pipeline for rapid analysis of targeted multi-pathogen genomic data

**DOI:** 10.1101/2024.06.28.601013

**Authors:** Richard Mayne, Shannah Secret, Cyndi Geoghegan, Amy Trebes, Kai Kean, Kaitlin Reid, Gu-Lung Lin, M. Azim Ansari, Mariateresa de Cesare, David Bonsall, Ivo Elliott, Paolo Piazza, Anthony Brown, James Bray, Julian C. Knight, Heli Harvala, Judith Breuer, Peter Simmonds, Rory J. Bowden, Tanya Golubchik

## Abstract

**Motivation:** Target enrichment strategies generate genomic data from multiple pathogens in a single process, greatly improving sensitivity over metagenomic sequencing and enabling cost-effective, high throughput surveillance and clinical applications. However, uptake by research and clinical laboratories is constrained by an absence of computational tools that are specifically designed for the analysis of multi-pathogen enrichment sequence data. Here we present the Castanet pipeline: an analysis pipeline for end-to-end processing and consensus sequence generation for use with multi-pathogen enrichment sequencing data. Castanet is designed to work with short-read data produced by existing targeted enrichment strategies, but can be readily deployed on any BAM file generated by another methodology. It is packaged with usability features, including graphical interface and installer script.

**Results:** In addition to genome reconstruction, Castanet reports method-specific metrics that enable quantification of capture efficiency, estimation of pathogen load, differentiation of low-level positives from contamination, and assessment of sequencing quality. Castanet can be used as a traditional end-to-end pipeline for consensus generation, but its strength lies in the ability to process a flexible, pre-defined set of pathogens of interest directly from multi-pathogen enrichment experiments. In our tests, Castanet consensus sequences were accurate reconstructions of reference sequences, including in instances where multiple strains of the same pathogen were present. Castanet performs effectively on standard laptop computers and can process the entire output of a 96-sample enrichment sequencing run (50M reads) using a single batch process command, in *<* 2 h.

**Availability and Implementation:** Source code freely available under GPL-3 license at https://github.com/MultipathogenGenomics/castanet, implemented in Python 3.10 and supported in Ubuntu Linux 22.04 and other Bash-like environments. The data for this study have been deposited in the European Nucleotide Archive (ENA) at EMBL-EBI under accession number PRJEB77004.

## Introduction

The popularity of metagenomics for sequencing multiple organisms from clinical samples has generated increased interest in strategies to enrich metagenomic libraries for organisms of interest, both to reduce per-sample cost and to improve sensitivity. Strategies to reduce unwanted sequence background are based on either depleting known host-derived sequences, such as human ribosomal genes, or on enriching targets of interest through amplification strategies. An increasingly popular choice is targeted enrichment by hybrid capture, also called ‘targeted sequencing’ or ‘bait capture’. Targeted sequencing involves the hybridisation of the metagenomic library with a potentially large panel of oligonucleotide probes, followed by amplification of the probe-bound sequences to ideally 100–1000*×* above the metagenomic background. This process has been shown to be unbiased at probe-target divergence of up to 20% (1), providing a convenient approach for detecting and sequencing even highly diverse pathogens such hepatitis C virus (1; 2), as well as for the analysis of within-host variation as has been done for respiratory syncytial virus (RSV) (3), human immunodeficiency virus (HIV) (4; 5) and SARS-CoV-2 (6). Target capture is seen use as a lower-cost, high-sensitivity screening method for SARS-CoV-2 variants not adequately covered by extant PCR primers (6) and is a potential successor to conventional amplicon bacterial 16s sequencing, as such methods greatly reduce the need for additional culture of difficult-to-cultivate organisms (7; 8). Encouraging progress has also been made in use for molecular profiling of tumours via targeted sequencing of cell-free tumour DNA to improve on conventional biopsy-based diagnosis (9).

A key feature of targeted enrichment is that the oligonucleotide panel can be customised to include organisms or genes of interest, forming an explicit set of hypotheses against which results can be assessed. This is in contrast with shotgun metagenomic sequencing, where the entire nucleic acid extract is sequenced, and expert judgement must be made about the potential clinical significance of every identified organism. An additional difficulty of metagenomic analysis is that most computational pipelines, such as QIIME, MetaPhlan and Kraken, (10; 11; 12), are designed to quantify and report relative abundance and/or species richness in the sample, with taxonomic classification as a key step, rather than being tuned to detect a potentially large set of organisms of particular interest. Abundance-based analysis methodologies are powerful when the full sample composition is of interest, e.g. in microbiome sequencing, as organism loads often may have environmental/clinical correlates and feed in to downstream analyses. They are not, however, optimal when the aim is to determine and quantify the amount of one or more specific organisms or loci, which is achieved more efficiently with target enrichment.

From a computational perspective, targeted sequence data has several attractive features that can be leveraged during analysis. First, the panel of targets can be pre-defined, making it possible to use established, highly efficient reference-based mapping strategies for sequence processing, with only minor modifications. Second, since a background level of (un-targeted) metagenomic reads remains in the sequenceable library, the difference in read depth between the targeted and untargeted sequences can be used to calculate efficiency of capture, and potentially distinguish true low-abundance positives from low-level background reads. Targeted sequencing combined with reference mapping also allows for read de-duplication, and therefore lends itself to quantitative analysis of input material, equivalent to estimating pathogen load directly from sequence data, as has been demonstrated for viral load in HIV (4), RSV (3; 13) and others (14). Although these features of targeted sequencing are likely appreciated within laboratories that use capture, wider uptake of targeted sequencing requires bioinformatics tools to support the specific features of the laboratory method.

All diagnostic sequencing workflows share some basic principles, including subtraction of unwanted host (e.g. human) reads and read trimming or filtering to remove sequencing artefact and low-quality sequences. Rapid development of laboratory methods has resulted in a need for cross-validation of multiple pipelines as part of an experimental design (15; 16; 17). Although data processing steps vary, analysis usually requires multiple steps that involve taxonomic classification and/or read quantification. Downstream genome assembly may then be conducted for each identified pathogen, either *de novo* (reference-free) or by mapping the reads to a specific reference sequence. In either case, analysis implies repetition of the same set of steps for every pathogen of interest. Analytical functions typically involve multiple tools for generating statistics from mapped read files using standard libraries e.g. samtools (18), and extracting meaningful sequence reconstructions (e.g. Gencore (19), shiver (20), ncov-19 (21)).

Here we present the Castanet pipeline (hereafter ‘Castanet’), an end-to-end, automated pipeline specifically designed for the analysis of targeted sequencing data. Our motivation was to create an efficient, easy-to-use, modifiable pipeline that enables rapid processing of a targeted sequencing experiment, and outputs all relevant metrics for downstream analysis. Castanet produces consensus sequences for targets identified in the data, along with summary statistics per target and per organism.

User input to Castanet consists of paired read files (fastq with or without compression) and a mapping reference (fasta file) containing one or more targets of interest, although an alternate endpoint exists for users to supply pre-mapped reads (BAM file) instead of the former.

Castanet is based on several open source tools, as well as additional methods for classification and quantification (Fig. 1) based on aggregation of reads by organism across multiple target sequences. This depends on an initial mapping to a collection of target sequences (the ‘mapping reference’). The reference will normally reflect the set of targets against which the capture probes were designed, although need not match precisely, as the selective amplification of specific targets enables prior assumption of sequence composition within a sample. Control strategies can be supported; for example, a reference sequence that contains both a targeted and an untargeted region can be used to estimate enrichment efficiency.

**Fig. 1.**
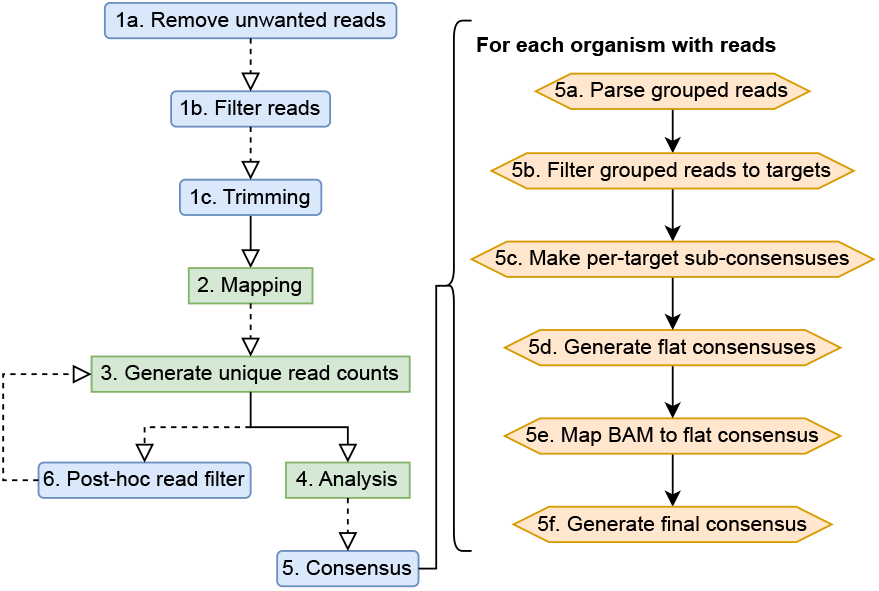
Castanet process flow. (Left) Castanet pipeline. Blue boxes are optional stages. (Right) Consensus generator algorithm.

Castanet’s analytical functions are species-agnostic and examine the comparative distribution of duplicated and deduplicated (unique) reads and their genomic positions. This allows for estimation of pathogen abundance (viral load), analysis of capture efficiency and efficient elimination of background reads. Aggregation of reads mapped to multiple targets to the level of individual organisms is achieved by interpretation of target nomenclature via natural language processing functions, according to defined rules. Statistics are then generated for every organism where reads have been detected, including coverage, number of targets/loci mapped, amplification rate and read proportions. Finally, consensus sequences are generated where coverage is sufficient, via an algorithm that generates ‘sub-consensuses’ for each target before they are flattened (taking majority bases from a partial alignment of reads and reference sequence) and reads are re-mapped to this as a reference. Crucially, this does not derive spans of consensus sequences from their references, which allows for more precise downstream analyses such as strain typing.

Castanet was originally written as an in-house research tool (14), implemented in Python 2 (https://github.com/tgolubch/castanet); this experimental version has been publicly available since 2019 and has been in extensive use within our own laboratories since 2018. To maximise the utility of the software for the bioinformatics community, we have now updated, optimised, fully automated and substantially expanded functionality to include consensus calling and utility functions. Aims for the Castanet application included:

- **User friendliness** Extensive bioinformatics experience is not required to install and run the software. Automated build scripts and a graphical user interface (GUI) are available, and command-line (CLI) entrypoints are available for expert users.
- **Automation** No manual curation is required to run Castanet other than ensuring the mapping reference is generated in a multi-fasta file with an appropriate sequence naming convention.
- **Performance** A paired read file of approx. 1 *×* 10^6^ reads may be analysed in under 5 minutes, on a consumer-grade laptop with a 16 thread 3.30 GHz processor and 32 Gb RAM.

Here we detail experiments validating Castanet and conclude by critically evaluating results in the context of the current gold standard methods.

## Materials and Methods

### Software

Castanet is written in Python 3.10 and is native to Ubuntu 22.04, but is also compatible with Bash-like environments including Mac. The application is hosted on a lightweight Uvicorn server behind a RESTful API composed in FastAPI (22) and may be deployed with a single command. The FastAPI framework has a GUI that may be accessed from the host’s browser, but may also be interacted with via the command line. All external dependencies may be installed via a shell script included in the repository. A full readme is included in the repository.

Individual pipeline stages (Fig. 1) are detailed below, with external dependencies listed in bold. The pipeline may be accessed via API endpoints as a single end-to-end run, a batch run that will iteratively fire end-to-end runs for all datasets within a single folder, or individual pipeline stages. Users may, for example, skip steps 1-4 if they already have a BAM file from a different source.

1. (Optional) Pre-process data:
  a. Taxonomically classify reads (call **Kraken2**).
  b. Filter out unwanted reads based on taxonomy, e.g. arising from the host genome.
  c. Trim adapters and remove low quality reads (call **Trimmomatic** (23)).
2. Map cleaned reads to reference sequences to produce a BAM file containing all mapped reads, including improper pairs (call **BWA-mem2** (24), **Samtools**).
3. Parse BAM file:
  a. Aggregate reads by target. Remove spurious matches falling below a user-defined threshold on fragment length (TLEN). Improperly paired reads are allowed if the paired match is to another variant of the same target, and is of sufficient mapped length.
  b. Generate total and unique (position-deduplicated) read counts.
  c. Temporarily cast reads grouped by target to separate BAM files.
4. Call analysis functions:
  a. Parse reference lengths, merge with position counts.
  b. Aggregate target sequences as organism-locus, via regular expressions. Organism-locus can be understood as a group of non-homologous regions that comprise the entirety of targeted sequence for a specific organism, e.g. multiple genes from a bacterial genome.
  c. Generate statistics for each organism-locus.
  d. Generate approximate coverage plots for each organism-locus.
5. Reconstruct consensus sequences (Fig. 1). Gaps in genomes with incomplete coverage are not filled by the reference. For each organism with reads:
  a. Parse all reads grouped by target from step 3.
  b. For each organism with reads, filter reads to each individual target, via collation from step 4.
  c. Generate per-target ‘sub-consensus’ sequences (**Samtools**).
  d. Create ‘flat consensus’: filter master BAM to organism-specific targets with coverage and depth exceeding user-specified thresholds and align sub-consensuses with relevant reference sequences, then generate a consensus (**Samtools, Mafft** (25)).
  e. Remap filtered master BAM from step 2 to flat consensus (**BWA-mem2**).
  f. Call final consensus on remapped flat consensus (**ViralConsensus** (26))
6. (Optional) Output of analysis (misassigned reads) used to filter input BAM file, re-call step 3 (**Samtools**).

### Experimental Evaluation

To evaluate Castanet, we used five datasets: two synthetic and three real (Table 1). To focus the testing strategy on what is a primary use of Castanet in our laboratory, all experiments focused on detection of viral agents. We used a mixture of data with unknown (set A) and known (set B) genomic content to compare Castanet’s utility in both ‘real-world’ and benchmark contexts.

**Table 1.** Dataset descriptions, excluding viral multiplex control. N targets: number of target sequences to map against in probe set.

| Expected Organism | Set | Genome length (bp) | N targets | N Samples | Method | Specimen type |
| --- | --- | --- | --- | --- | --- | --- |
| Viral multiplex control | A.1 | Various | 5 | 5 | Target enrichment | Reference material |
| Negative Plasma | A.2 | N/A | N/A | 5 | Target enrichment | Plasma |
| Marburg virus | B.1.1 | $1.9 \times 10^4$ | 5 | 3 | Synthetic | N/A |
| HCV-1A, 2A, 2B | B.1.2 | $\sim 9.4 \times 10^3$ | 3 | 3 | Synthetic | N/A |
| RSV-A, B | B.2.1 | $\sim 1.5 \times 10^4$ | 12 | 17 | Target enrichment | Nasopharyngeal swab |
| HIV-1 | B.2.2 | $\sim 9.7 \times 10^3$ | 9 | 21 | MTA-Seq | Whole blood |

For set A (ENA PRJEB77004), Castanet was used to analyse a dataset generated in-house (SI Document 1, S1) via target capture as a simulated clinical application and included:

A.1 Viral multiplex reference (National Institute for Biological Standards and Control, London, United Kingdom). Pre-mixed material from 25 viruses of human and bovine origin, of which the following were quantified in International Units: Parechovirus (HPeV), Human Herpes Virus 4 (HHV4, more commonly known as Epstein-Barr virus, EBV), Human Herpes Virus 5 (HHV5, more commonly known as cytomegalovirus, CMV). This control was included as a dilution series to support estimation of viral load.
A.2 Five pooled clinical plasma samples from healthy donors, confirmed as PCR-negative for multiple infectious agents including HIV, Hepatitis B, C and E. These were included as being representative of negative samples in a screening context, that nevertheless could contain low numbers of reads for adventitious, non-pathogenic viruses such as torquetenovirus (TTV). Set B included both simulated (set B.1, (SI Document 1, S2)) and publicly-available data (set B.2). These included:
B.1.1 Marburg virus is a filamentous negative-sense ssRNA virus with three distinct clusters where intra-species nucleotide divergence is *<* 20% (27). This represents a straightforward use case where retrieval of the entire genome is feasible through use of a small number of probes.
B.1.2 Hepatitis C virus (HCV), equal mixture of reads from types 1a, 2a and 2b genotypes. This simulates a complex but relatively common clinical issue where multiple viral subtypes or quasi-species may co-exist within a patient (28). As a short positive-sense ssRNA virus, full genome coverage may easily be achieved with a small number of probes.
B.2.1 RSV. 17 clinical respiratory samples extracted from nasal swabs, in infants infected with this linear negative-sense ssRNA virus (ENA ERR10812874–86) (13). Samples were collected from UK and Spanish infants aged less than one year during the 2017–18 RSV season for the purpose of investigating virus subtype distribution and differential gene expression in this outbreak. The original study identified and published several diverse, novel isolates through a curated workflow using Shiver (20); the current study did not omit the four samples that the original study’s authors were not able to extract reference sequences from.
B.2.2 HIV-1. 21 clinical blood samples from individuals known to be HIV-1 positive, collected as part of a pan-European project (ENA ERR732065–90) for the purpose of providing open source data for HIV researchers (20; 29). HIV are retroviruses possessing duplicate copies of positive-sense ssRNA genomes whose capability for rapid mutation frustrates efforts to treat and study it (30), hence reconstruction of HIV genomes from NGS data is challenging because an individual with HIV is likely to be host to multiple quasi-species of the virus with significant intra- and inter-host divergence. HIV-1 sequences were pre-determined using a simple pipeline of Iterative Virus Assembler (IVA) (29) followed by Shiver, using default settings for both.

Descriptive statistics were collected including number of mapped reads, mean depth, coverage at depths 1 and 10, mean amplification rate (MAR, number of reads ÷ number of deduplicated reads at each position supported by reads) and consensus sequence quality were recorded from Castanet’s default output. Consensus quality was measured by comparing Castanet’s output with published reference sequences, using BLASTn (query coverage and percentage identity, hits where E values *<* 0.01) and Mash scores (31). The rationale for adopting two metrics was that Mash is a more sensitive tool and hence provides more information in instances where BLAST scores are very high.

We used an in-house mapping reference of 3682 sequences against a range of clinically-relevant viral, fungal, parasitic and bacterial targets, including ribosomal multi-locus (rMLST) (32) targets for the latter, in all experiments. Castanet’s default settings were used throughout.

### Hardware

All experiments were conducted on a laptop with 16 hyperthreading-enabled cores (i9-11980HK, 3.30 GHz), 32 Gb DDR4 RAM and M2.SSD with read/write speed of 6 *×* 10^6^ Mb s^−1^, running Windows Subsystems Linux Ubuntu 22.04 LTS. Experiment run time was benchmarked in all experiments.

## Results

The average run time for a single sample across all experiments conducted between sample sets A–B was 7.28 min (mean read depth 5.2 *×* 10^5^), the longest run in 31.0 min which comprised over 3 *×* 10^6^ reads and involved generating consensus sequences for 4 organisms. Compute time scaled linearly (SI Document 1, S3) with read count; no experiments saturated the computer’s memory. The most costly functions were consensus generation and analysis (see section 2.1, steps 6–7), each comprising approximately 40% of run time.

For detected organisms of interest, plots of read depth across the span of each genome for both deduplicated (unique) and total reads were generated automatically by Castanet (Fig. 2a–b). These were used to assess coverage and, in the case of pre-quantified controls, estimate viral loads (Fig. 2c) using linear least squares regression of log_10_ of deduplicated reads reported by Castanet versus log_10_ of known laboratory-reported viral load from qPCR experiments, in the three viruses from set A.1 (HPeV, EBV, CMV).

**Fig. 2.**
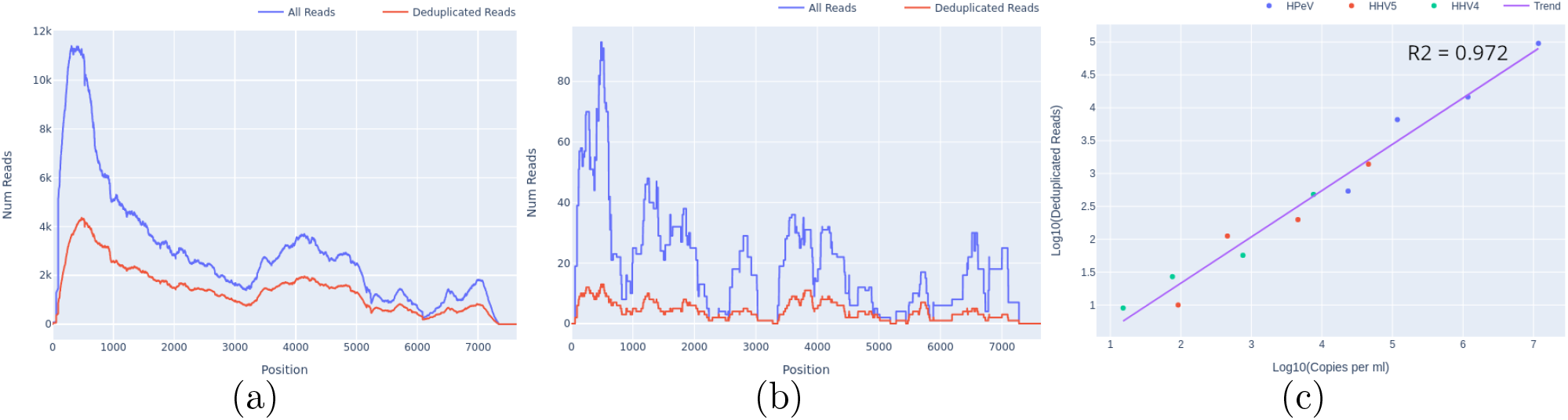
Castanet output for set A.1, at several dilutions. (a) Read depth graph showing total and deduplicated (unique) reads for HPeV, undiluted, viral load 9.6 *×* 10^4^ copies per ml. (b) Read depth, HPeV, 1:500 dilution, 5.3 *×* 10^2^ copies per ml. (c) Log10 deduplicated reads versus Log10 copies per ml for HPeV, HHV4 and HHV5 at four different dilutions (1, 1:10, 1:100, 1:500), with linear regression line.

A significant benefit of targeted sequencing is that signal of enrichment may be useful for distinguishing true low abundance positives from the background sequences that are a ubiquitous feature of all sequencing experiments (33). In our experience, genome coverage plots of low-abundance true positives that have been successfully sequenced with target enrichment are expected to show a characteristic “city-block” appearance, with a series of highly amplified fragments across the targeted regions, which result from capture and amplification of a small number of template fragments in the sample. We observed such patterns in both controls at known low titres (Fig. 2b) and clinical samples where low viral loads were anticipated, e.g. commensal viruses such as TTV (Fig. 3a). In contrast, background sequence noise, such as index misassignment or carryover from other samples in the batch, will have very few or no duplicated reads at the targeted region, with no evidence of amplification which is what we observed (Fig. 3b). It is notable that the use of amplification intensity to distinguish low-abundance positives from background noise is method-specific; it is well-suited to targeted enrichment protocols, and generally does not apply to shotgun metagenomics, allowing for certain edge cases, e.g. index misassignment, contamination from another library. Consequently, when untargeted shotgun data is analysed with Castanet, the plots can be expected to show very little sequence duplication (SI Document 1, S4).

**Fig. 3.**
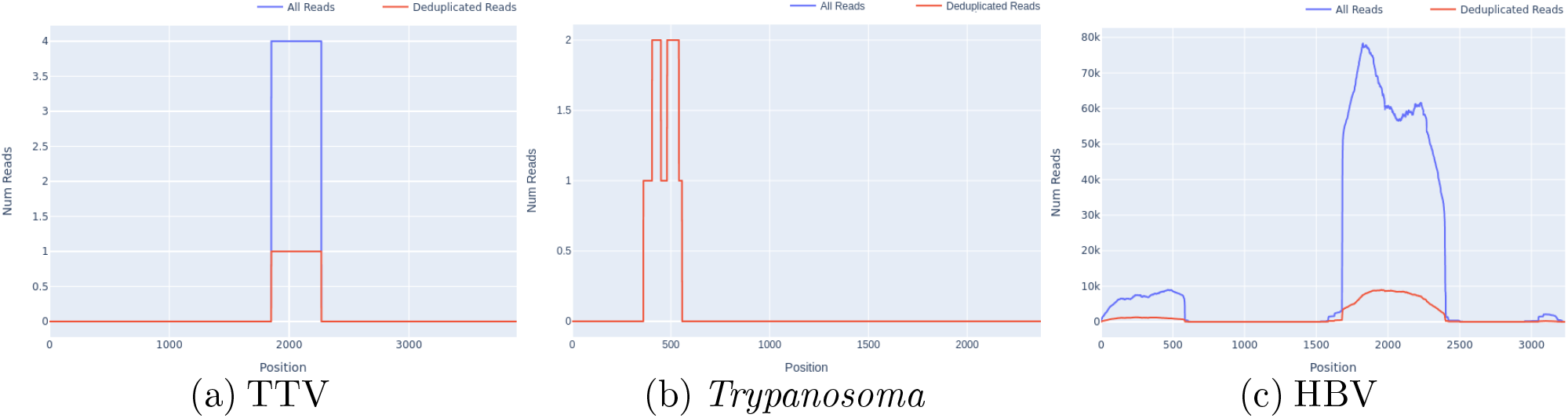
Appearance of presumptive low positives, false positives and contamination in read depth plots from set A.2. (a) Reads for TTV, with high amplification rate across the region with reads (4.0), across a partial region of the genome. (b) *Trypanosoma* spp. reads at the ribosomal 18s locus, with no amplification and incomplete coverage. (c) HBV reads with incomplete coverage and significant amplification (MAR 5.82, S.D. 3.37).

A common source of difficulties with standard taxonomic classification tools for metagenomics data, which quantify taxa on the basis of read identity, is that reads may originate from a high-abundance contaminating source, such as pre-amplified PCR product. In standard metagenomics processing, such contaminated samples will yield extremely high numbers of reads matching a given taxon and will be considered high-confidence positives. However, using reference mapping, contamination of the sample by e.g. pre-amplified (PCR) product is readily detectable as unusually high depth across a defined proportion of the targeted region, with no evidence of sequence across the rest of the targets for the same organism. An example of this type of contamination is shown in Fig. 3c, where an HBV sample was accidentally contaminated by a small amount of amplified product from a diagnostic PCR in the originating laboratory, seen as a discrete region of high depth at the targeted genomic region, and a smaller region of cross-mapping of some reads to an imperfect genomic repeat of the target. Castanet uses mapping to make this analysis intuitive.

Another crucial process in analysing a target enrichment experiment is estimating the efficiency of capture, as many downstream applications rely on amplification of target sequences being sufficient to achieve adequate coverage and read depth to properly represent sample diversity. This may be achieved with Castanet’s output in several ways, including by comparing the mean sequencing depth between targeted and non-targeted sequences. Non-targeted sequences are part of the uncaptured (metagenomic) background, and hence would be expected to have little-to-no evidence of read duplication, whereas targeted sequences should have significant amplification (100–1000*×* the background) in cases where capture has been successful. In a representative example from set A.2, we found the read depth to be ∼ 130*×* that of the metagenomic background when comparing read depth of the whole human mitochondrial genome (Fig. 4a) with cytochrome C oxidase I (COX-1) (Fig. 4b), where the latter was the only targeted region in the former.

**Fig. 4.**
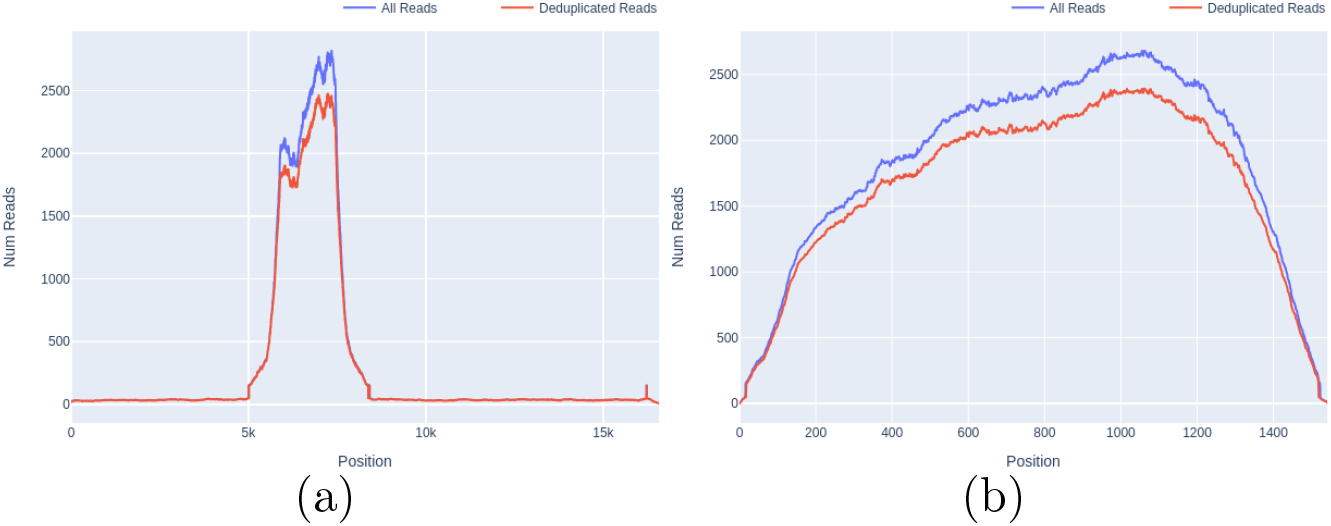
Comparison of read depth between untargeted and targeted sequences, in a screened pooled human plasma sample (set A.2). (a) Whole human mitochondrial gene, median depth 16 (excluding COX-1 region). (b) Targeted sequence, mitochondrial COX-1, median depth 2109.

A primary novelty of Castanet is its aggregation of reads to the correct organism, which it did correctly in all experiments from set B (Tab. 2). For consensus reconstruction, counts as low as 1 *×* 10^4^ were sufficient to recover the vast majority of genomic content in all experiments. For detection of organisms where a complete genome is not required, a significantly lower read number (approximately 1 *×* 10^2^) was sufficient.

**Table 2.** Summary statistics for Castanet validation on set B data. 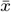: mean; parentheses indicate standard deviation on set B.2 (real) datasets. Con Rec: percentage of samples where consensus was recovered; N reads: number of reads on target; Depth: read depth; Amp rate: amplification rate (n reads *÷* n reads deduplicated); Cov: percentage coverage with depth *>* 1 / *>* 10; Con QC: consensus BLASTn query coverage versus reference sequence; Con PcID: consensus BLASTn percentage identity versus reference sequence; Con MH: consensus Mash score verses reference sequence.

|  | Marburg Virus |  |  | HCV-1A/2A/2B |  |  | HIV-1 | RSV-A, B |
| --- | --- | --- | --- | --- | --- | --- | --- | --- |
| | $1 \times 10^4$ | $1 \times 10^5$ | $1 \times 10^6$ | $3 \times 10^4$ | $3 \times 10^5$ | $3 \times 10^6$ | | |
| n | 10 | 10 | 10 | 10 | 10 | 10 | 21 | 17 |
| $\bar{x}$ Run time | 0.5 | 1.0 | 10 | 1.2 | 3.1 | 29 | 6.5 (4.7) | 8.5 (9.0) |
| Con Rec (%) | 100 | 100 | 100 | 100 | 100 | 100 | 100 | 76 |
| $\bar{x}$ N reads | $0.9 \times 10^4$ | $1.0 \times 10^5$ | $1.0 \times 10^6$ | $3.0 \times 10^4$ | $3.0 \times 10^5$ | $3.0 \times 10^6$ | $8.2 \times 10^4$ ($1.1 \times 10^4$) | $7.0 \times 10^4$ ($3.1 \times 10^4$) |
| $\bar{x}$ Depth | $2.6 \times 10^2$ | $2.6 \times 10^3$ | $2.6 \times 10^4$ | $1.5 \times 10^3$ | $1.5 \times 10^4$ | $1.5 \times 10^6$ | $2.6 \times 10^4$ ($1.7 \times 10^4$) | $1.4 \times 10^4$ ($1.9 \times 10^4$) |
| $\bar{x}$ Amp rate | 1.00 | 1.02 | 1.16 | 1.00 | 1.03 | 1.33 | 3.35 (1.39) | 2.98 (1.93) |
| $\bar{x}$ Cov 1 (%) | 99.9 | 99.9 | 99.9 | 98.1 | 98.2 | 98.2 | 98.32 (0.55) | 96.1 (6.71) |
| $\bar{x}$ Cov 10 (%) | 99.8 | 99.9 | 99.9 | 98.1 | 98.2 | 98.2 | 94.83 (7.26) | 83.7 (26.6) |
| $\bar{x}$ Con QC (%) | 90.0 | 99.0 | 99.1 | 99.0 | 100 | 100 | 92.3 (2.86) | 98.3 (1.55) |
| $\bar{x}$ Con PcID (%) | 99.9 | 99.9 | 99.9 | 99.9 | 100 | 100 | 98.6 (1.07) | 100 (0.02) |
| $\bar{x}$ Con MH (%) | 84.2 | 95.4 | 95.6 | 99.7 | 100 | 100 | 63.2 (16.1) | 98.7 (1.89) |

The quality of consensus sequences, measured via BLASTn and Mash scores comparing Castanet consensuses to reference sequences, increased to peak values exceeding 99% (BLASTn QC/PcID) and 95% (Mash) at read counts exceeding 1 *×* 10^5^ in set B.1. We were able to differentiate between and generate consensus sequences that approached 100% identity with reference sequences, for all three HCV subtypes in set B.1.2. Reads on target (i.e. mapped to the reference) in the order of 10^4^ were retrieved in both B.2 datasets, which was sufficient to generate coverage exceeding 83% in all cases, or 94% if excluding the four insufficient RSV samples. These four RSV samples had much lower mean coverage (mean 85.5%, std. d. 6.20% and mean 45.0%, st. d. 25.4% at depth 1 and 10, respectively) and MAR than other samples, although consensus sequences with sufficient coverage to support strain typing were generated for all, where this had not been achieved previously.

## Discussion

Castanet is designed to provide outputs of greatest interest to those running targeted multi-organism sequencing experiments, but is broadly applicable to sequence data generated using other methods. Reporting of statistics such as coverage and depth are essential in any sequencing workflow, as these indicate the relative success of the experiment and whether downstream analyses such as variant discovery will be possible. Castanet reports additional statistics such as MAR (an amplification of 1 indicates poor enrichment, or no enrichment or contamination): we used this in set B.2.1 experiments to demonstrate correlation of failure to recover a consensus genome with low amplification, which could result from experimental factors such as sample insufficiency. The full Castanet analysis output (SI Spreadsheet 1, Sheet 1) encompasses 51 statistics which allow for flexible characterisation of experimental data depending on experimental design.

Although we focused our reporting on the most abundant organisms in the samples described here, a range of other organisms were detected in the majority of B.2.1 samples including expected viral and bacterial species (human rhinovirus, *Moraxella catarrhalis, Haemophilus influenzae* among others) (SI Spreadsheet 1, Sheet 2). This highlights the utility of Castanet for rapid and comprehensive analysis of target capture data to characterise a wide variety of organisms in a single workflow which could be applied to diagnostic microbiology settings.

The quantitative relationship between deduplicated read counts and a sample’s viral load, described extensively in previous publications (13; 4; 6)), further illustrates how Castanet’s output may feasibly be used in conjunction with target capture methods to replace PCR testing in a diagnostic setting, provided sufficient quality control and calibration provided for as part of the workflow.

Contamination is a recognised risk in all high-throughput sequencing workflows, with multiple potential sources for it to be introduced into a library. Deriving the consensus sequence directly from initial analysis can help troubleshoot the likely source of contamination, e.g. in the case we examined (Fig. 3a), the contaminant was traced to PCR amplicon from a neighbouring laboratory source. We have found that a coverage proportion at minimum depth 2 of ∼ 10—15% to be a good indicator of likely true presence in problematic samples with low read depth, and *>* 30% is usually confirmatory, although these precise thresholds will differ between laboratories, methods and target organisms, so should not be considered without reference to the amplification rate. This highlights another justification for the use of target capture methodologies in high-throughput settings when leveraged via Castanet.

Users may evaluate coverage per-target and fine-tune consensus generation parameters where necessary, using the automatically-generated consensus alignment plots (SI Document 1, S5). This is relevant in cases such as set B.1.2, as it was known *a priori* that three HCV variants were present. Default Castanet behaviour would aggregate all reads to “Hepatitis C” and generate a consensus sequence represented by the majority subtype. There are multiple options for extracting more precise typing from Castanet, such as modification of mapping reference nomenclature, use of a separate mapping reference file where aggregation is allowed only at the level of subtypes, or extracting the sub-consensuses from the consensus module’s output. It may be sufficient for many workflows, however, to merely confirm the presence of reads against a specific organism with Castanet and use the consensus sequence to compare bulk genomic data between samples before doing downstream strain typing.

All consensus quality scores were lower in set B.2.2 experiments (against set B.2.1, consensus query coverage 92.3 vs 98.3%, percent identity 98.6 vs 100%, Mash score 63.2 vs 98.7%), as would be expected for viruses such as HIV-1 that form diverse intra-host populations, as alignment involves taking majority values across all sub-consensuses. In such cases it is possible to extend Castanet to add a species-specific workflow once the correct pathogen has been identified. HIV-1 data used for benchmarking in this study were generated with MTA-seq, which highlights the utility of the Castanet pipeline for different NGS techniques.

Comparisons of Castanet with the software used to generate reference sequences are included here to guide users towards the appropriate choice of analytical pipeline. In the case of our most difficult example, HIV, shiver was designed for optimal reconstruction of HIV genomes from highly diverse clinical samples by regenerating missing regions from the closest-matching reference sequence from a user-defined library, or from *de novo*-assembled contigs. This requires a well-curated database of reference sequences and possibly expert input to evaluate the quality of alignments before contigs are flattened with the references. Castanet output is suitable for further processing with shiver, but we chose not to implement additional specific specific workflows within the tool itself.

We have presented Castanet’s method of aggregating reads to specific targets at the level of organisms as a feature, but it is pertinent to note two limitations. First, the naming conventions of the user’s probe file must be kept consistent to ensure organism-level aggregation; three regular expressions are provided in Castanet’s documentation to guide users, and our future work includes further refinement of this step. Second, for consensus reconstruction, we assume genome collinearity with references, sample colonisation by a meaningful majority quasi-species per organism, and a theoretical coverage of 100% across the targeted regions per organism. In cases where only part of an organism’s genome is targeted, we recommend that the user defines the reference sequences appropriately to match the targeted regions, or sets the consensus parameters in line with a reduced requirement for genome coverage. This is true particularly where larger genomes such as those of bacteria or fungi may not be fully represented in a user’s target enrichment panel, and thus Castanet would be unable to recover a complete consensus genome in these instances. In general, the reference should be provided in accordance with the design of the experiment. Multipartite genomes may be processed into a linear approximation through concatenation in order of segment size, or retained as separate segments. Future work includes expanded support for multi-locus targets and segmented viral genomes.

Targeted sequencing is well-suited to diverse applications in both research and clinical practice, including pathogen genomics and transcriptomics. The capacity of large diagnostic laboratories to perform sequencing has increased greatly over recent years. As costs continue to fall, targeted sequencing is likely to supersede methods such as 16s sequencing to become part of the routine diagnostic armoury in clinical settings as combined diagnostic and surveillance testing. The Castanet pipeline facilitates bioinformatic analysis to take place outside research facilities, as it addresses the lack of open-source, end-to-end tools designed specifically to exploit targeted sequencing data and hence supports wider adoption of these technologies. Our analysis here has demonstrated how the software may be used to support a wide variety of applications as well as evaluate run performance and troubleshoot laboratory issues, in a scalable and reproducible manner.

## Acknowledegements

RM, SG, KK, KR, HH, JB, PS and TG are supported by funding from the UK National Institutes for Health Research (NIHR) [grant number NIHR203338]. TG is supported by an Investigator Grant from the National Health and Medical Research Council, Australia (NHMRC) [grant number GNT2025445]. JB is supported by the UCL/UCLH NIHR BRC. The authors are grateful to Prof. Martin Maiden for his guidance in implementing rMLST gene compatibility of our capture oligos and mapping reference.

