## Supplementary data, document 1 for "Castanet: a pipeline for rapid analysis of targeted multi-pathogen genomic data"

### S1. In-house NGS methods

DNA was extracted from pooled screened human plasma and a supplementary multi-blood-borne pathogen control sample (Zymo Research, CA, USA) and prepared for target capture using a bespoke panel of oligonucleotide probes (Twist Bioscience, CA, USA), according to the manufacturer's protocols. Sequencing was conducted on an Illumina MiSeq system to generate PE150 reads. Samples were sourced from the United Kingdom's NHS Blood and Transplant (NHSBT) as part of a project evaluating NGS methods for microbiology screening of transfusion and transplant donors red with internal approval of the project.

### S2. Synthetic read generation strategy

Synthetic sets were generated red using an NGS whole genome simulator, DWGSIM 0.1.14 (https://github.com/nh13/DWGSIM), at $1 \times 10^4$, $1 \times 10^5$ and $1 \times 10^6$ read depths in 250 bp pairs, using a constant error rate of 2.5%, chosen as an overestimation of mean error rates from modern sequencers.

### S3. Castanet benchmarking

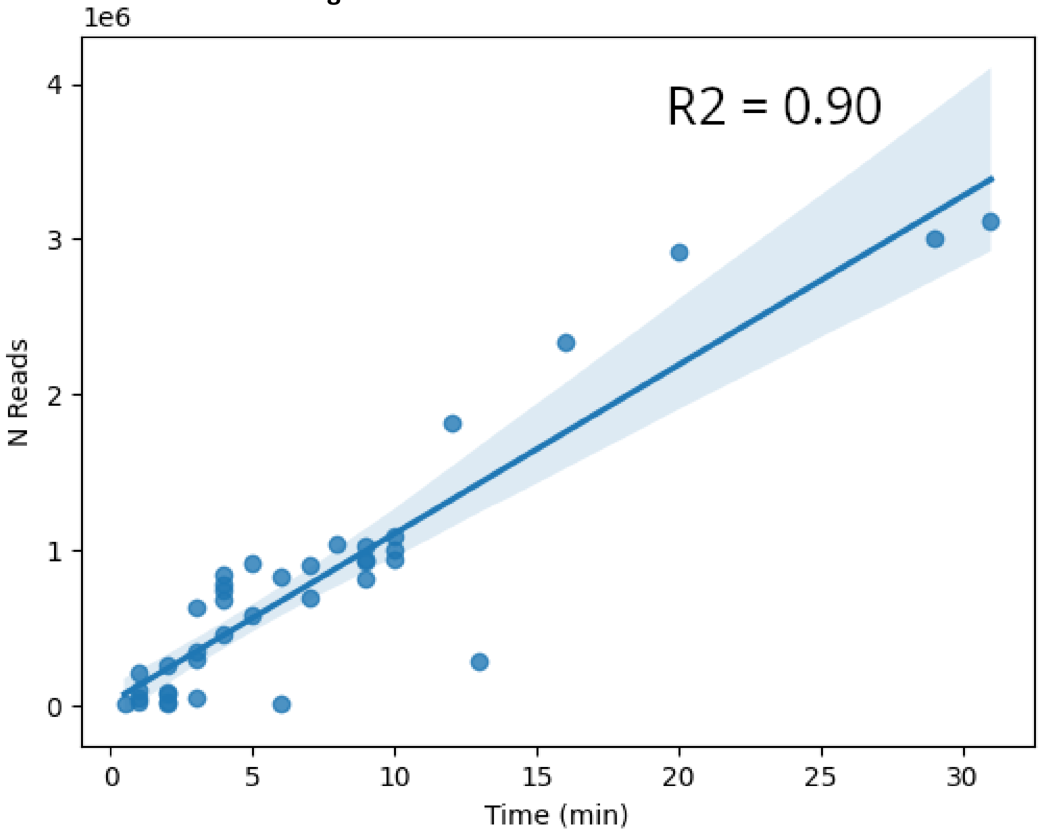


Figure S3.1. Castanet performance benchmarking. Compute time versus read count, with linear trendline and 95% confidence interval.

### S4. Castanet applied to metagenomics data

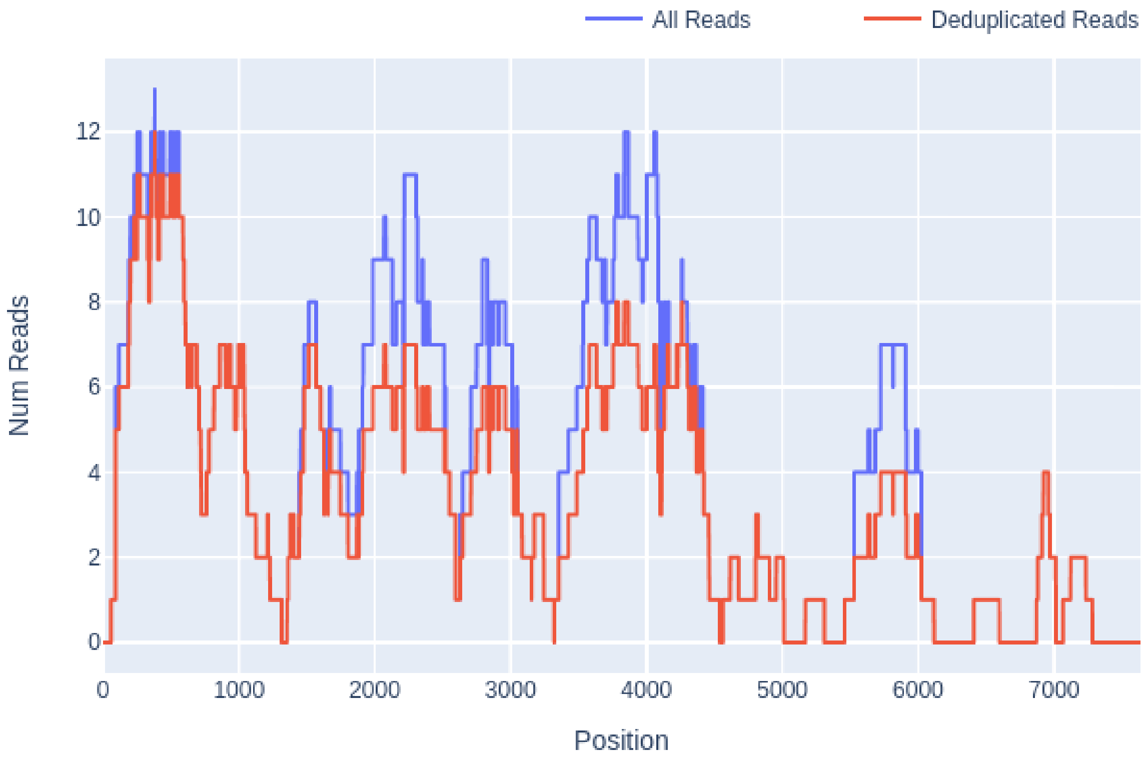


Figure S4.1. Read depth (total and deduplicated), HPeV, sample set A.1, 1:10 dilution, metagenomics sequencing.

A metagenomics (shotgun) sequencing experiment on dataset A was run in parallel to target capture experiments, using pooled pre-capture libraries from in-house sequencing on dataset. Sequencing was performed on a NovaSeq X using a partial lane, sharing 100 Gb reads between 96 samples; otherwise, sample preparation and analysis was as in the experiments described in the main text. Coverage plots showed significantly lower levels of sequence deduplication (Fig. S5.1) than in target capture run (Fig. 2a, main text).

### S5. Castanet alignment plots: example and description.

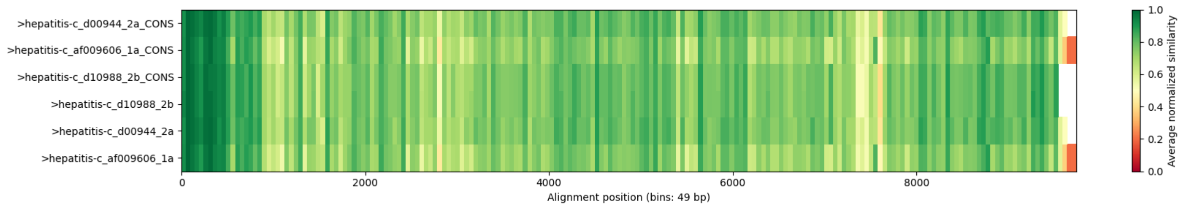


Figure S5.1. Castanet alignment plot, showing binned average normalised similarities between mapping references (">hepatitis-c_...") and sub-consensuses (">hepatitis-c_... _CONS").

Castanet's consensus sequence generator creates several visualisations to aid users in interpreting their results. In addition to consensus coverage and identity plots (data not shown), it also generates a heatmap showing the average normalised similarity between binned sequences from both mapping references and sub-consensuses (Fig. S2.1). This function is based on the Python library Biotite (https://www.biotite-python.org/) and may be used to evaluate the appropriateness of using Castanet to aggregate reads to a single mapping reference species in complex cases, such as in the example above, where diverse populations of quasi-species are anticipated to be present in a sample. The example above shows high similarity of each sub-consensus with their respective mapping references for three types of Hepatitis C virus, indicating the likely presence of all three in the sample.
